# The strength of selection for costly toxin production increases with growth rate

**DOI:** 10.1101/2024.12.09.627242

**Authors:** Ave T. Bisesi, Jeremy M. Chacón, Michael J. Smanski, William R. Harcombe

## Abstract

Microbes adopt a diversity of strategies to successfully compete with coexisting strains for space and resources. One common strategy is the production of toxic compounds to inhibit competitors, but the strength and direction of selection for this strategy varies depending on the environment. In particular, existing theoretical and experimental evidence suggests growth in spatially-structured environments makes toxin production more beneficial because competitive interactions are localized. Because higher growth rates increase the localization of resource competition in a structured environment, theory predicts that toxin production should be especially beneficial under these conditions. We tested this hypothesis by developing a genome-scale metabolic modeling approach and complementing it with comparative genomics to investigate the impact of growth rate on selection for costly toxin production. Our modeling approach expands the current abilities of the dynamic flux balance analysis platform COMETS to incorporate signaling and toxin production. Using this capability, we find that our modeling framework predicts that the strength of selection for toxin production increases as growth rate increases. This finding is supported by comparative genomics analyses that include diverse microbial species. Our work emphasizes that toxin production is more likely to be maintained in rapidly-growing, spatially-structured communities, thus improving our ability to manage critical microbial communities and informing natural product discovery.

## INTRODUCTION

Most microbial species exist in complex and dynamic communities where they must compete for limited nutrients and space [1–2]. Many microbes - particularly those from soil environments - deal with this competition by synthesizing a range of energetically costly antimicrobial compounds that can inhibit nearby sensitive species to gain a competitive advantage [2]. By reducing the density of toxin-sensitive species, toxin-producing strains can play an important role in structuring microbial communities over ecological and evolutionary timescales, with consequences for community function and ecosystem services. Additionally, bacterial toxins and related natural products can range from strain-specific bacteriocins to broad-spectrum antibiotics [3–4], and many of these secondary metabolites have diverse and beneficial applications such as herbicides or anticancer agents [5]. Understanding the conditions under which toxin production is beneficial is key to effectively managing microbial communities and advancing natural product discovery.

Theory predicts that the environment type and the localization of interactions are essential determinants of selection for toxin production. For example, selection against the production of diffusive toxins is well-documented in liquid (well-mixed) environments [2]. In such environments, interactions between microbes are global, such that all individuals have roughly equal access to resources and exposure to toxins. Under these conditions, toxins often remain too dilute to meaningfully inhibit competitors, and toxin producers are not able to invade established susceptible populations [2]. Similarly, when toxins are a global resource, resistant cheaters - individuals who are neither sensitive to the toxin nor produce it, therefore benefiting from production without paying an energetic cost - can easily outcompete toxin producers, selecting against production [6]. The localization of interactions afforded by spatial structure can offset both problems [3]. When strains grow in a spatially-structured environment, toxins will generally remain more concentrated near the site of production and kill any sensitive local competitors, allowing producers to make use of resources that would otherwise be assimilated by susceptible microbes [7–8]. Likewise, when resistant cheaters are present, they must typically be within a certain physical proximity to producers to benefit, protecting toxin producers from being outcompeted by cheaters [9]. By increasing interaction localization over well-mixed environments, spatially-structured environments set the stage for the success of toxin producers.

However, toxin production is not universally maintained in structured environments. For example, the strength of selection for production can decrease when toxin producers secrete high concentrations of antimicrobial compounds, because at high concentrations these compounds diffuse to more distant colonies and benefit resistant cheaters that are not growing nearby [9].

This underscores that toxin production - despite its observed importance in many microbial communities - can be challenging to select for and depends fundamentally on the scale of competitive interactions. The extent of interaction localization in spatially-structured environments is determined by a number of environmental and biophysical variables, including microbial growth rate [10–11]. We have previously demonstrated that growth rate impacts the scale of resource competition, such that interactions are more localized when bacteria grow quickly relative to metabolite diffusion [10]. This is because isolated microbes can rapidly use local nutrients before they diffuse to distant colonies and therefore grow to larger population sizes [10]. In contrast, when bacterial growth rates are slow relative to resource diffusion, some resources that are local at the beginning of growth will diffuse away from colonies before they can be used. As a result, when growth rates are low, interactions become more global - more closely resembling a well-mixed environment - and colonies grow to similar sizes [10]. For this reason, in spatially-structured environments, we hypothesized that toxin production should be more favorable as growth rates increase because interactions will become increasingly localized.

We tested these hypotheses with a combination of simulation modeling and comparative genomics. We augmented the capabilities of the COMETS modeling platform [12], which can predict microbial community dynamics using intracellular metabolic mechanisms as captured by genome-scale metabolic models. COMETS relies on dynamic flux balance analysis (dFBA) to predict the metabolic activity of species as a function of resource availability. While the platform tracks microbial growth and metabolite concentrations over time, we added the ability for chemical compounds to act as signals that modify the maximal growth rate, death rate, or bounds on metabolic reactions of a metabolic model. This capability makes it possible for COMETS to quantitatively predict the dynamics of microbial communities that include signals or toxins.

Using this platform, we simulated communities of bacteria composed of toxin producers, toxin-resistant cheaters, and susceptible competitors, altering bacterial growth rate to examine its impact on ecological selection for toxin producers [6, 9]. Finally, we curated two genomic datasets and tested the relationship between microbial growth rate and the presence of secondary metabolite gene clusters in a variety of bacterial species. We chose one dataset to leverage *in vitro* measurements of maximum growth rates across diverse microbial species, in addition to examining the relationship between predicted growth rate and toxin production in *Streptomyces*, a genus of soil bacteria well-known for producing a range of secondary metabolites [13–14].

Both our modeling and genomics approaches are consistent with the prediction that increasing growth rate generally strengthens selection for toxin production. These findings improve our ability to predict and manage microbial communities in spatially-structured environments.

## MATERIALS AND METHODS

### Modeling toxin generation and response

Toxin generation and response were integrated into the dynamic flux balance analysis modeling platform COMETS [12]. Generic microbial models were generated for three strains: a toxin producer, a toxin-resistant cheater, and a susceptible competitor. All three models were assigned three core reactions: carbon exchange (extracellular to intracellular), carbon transport (within the cell), and biomass generation. For resistant cheaters, no additional reactions were added to the metabolic model. For simplicity, we modeled toxin production occurring simultaneously with growth.

Production was implemented in producer models by having the model produce an extracellular metabolite in direct proportion to the flux through the biomass reaction. In other words, toxin production was encoded as a product in the biomass reaction, and an exchange reaction was included so that all produced toxin ended up in the extracellular environment. This meant that producers had two metabolic reactions in addition to the core three: toxin exchange (intracellular to extracellular) and toxin transport (within the cell). Toxin production could also be energetically costly, reflecting a fitness trade-off for the phenotype, such that a percentage of carbon went to biomass while the remainder was sequestered into toxin. The biomass equation of a toxin producer therefore became: *-(1 + cost_of_production) **carbon_c** → mmol_toxin_per_gram_cells * (1 + cost_of_production) **toxin_c*** where **carbon_c** and **toxin_c** are metabolites and all other terms represent reaction coefficients.

For susceptible competitors, we reduced the allowable upper bound on flux through the biomass reaction as a function of the extracellular toxin concentration. Our modeling framework can accommodate different functional relationships between toxin concentration and flux through the susceptible biomass reaction. A variety of parameterizing tests suggested that the functional relationship generally did not qualitatively change our simulation results. For this reason, we used a bounded linear relationship for all simulations, such that toxin concentration linearly reduced the upper bound of susceptible biomass flux and growth rates could not become negative due to toxin. Susceptibles therefore achieved a maximum growth rate when zero toxin was present (i.e. no change in upper bound) and then decreased linearly until there was zero possible growth when the toxin concentration reached 15 mmol (i.e. upper bound on growth = 0). We chose this constraint because, regardless of differences in maximum growth rate between conditions, all susceptibles would face the same relative effect of a given toxin concentration in terms of a percentage decrease in growth rate. Models of susceptible competitors had four total metabolic reactions, with toxin exchange (extracellular to intracellular) included alongside the core metabolic reactions. Resistant cheaters and susceptible models were functionally equivalent, except that resistant cheaters did not possess metabolic reactions involving toxin response.

We varied the maximum growth rate of all three models by varying the maximum carbon uptake rate. Because 1 gram of model biomass was produced per 1 mmol of carbon (except for simulations where we included a biomass cost of toxin production), we could limit maximum growth to 1 hr^-1^ by setting a 1 mmol/gDW/hr uptake bound on carbon exchange. Additionally, we did not remove toxins from the simulation, so they built-up and diffused in the environment like other waste products. Additional technical details of the implementation, and a guide to using it with COMETSPy, are available at https://bisesi.github.io/Toxins-Growth-Rate/.

### Toxin simulation model dataset curation and analysis

We completed batch-culture simulations in either well-mixed or spatially-structured environments, meaning the environment was seeded with nutrients once and depleted over the duration of the simulation. To evaluate the ability of a toxin producer to invade an existing susceptible population and outgrow a resistant cheater, we seeded initial susceptible densities at 80% of the total population (8e-9 gDW total biomass, with each of 40 susceptible colonies initialized at a biomass of 2e-10 gDW), with resistant cheaters and toxin producers each starting at 10% of the total population (1e-9 gDW total biomass, with each of 5 resistant and 5 producer colonies initialized at a biomass of 2e-10 gDW). We then varied the growth rate of all three strains from 0.125 to 1 hr^-1^, representative of a realistic range of growth rates [13], with the toxin producer paying a growth rate cost of 1% (i.e. on the same amount of carbon, producers can reach 99% of the biomass achieved by susceptibles or resistant cheaters). The total resource abundance and initial colony biomass inoculation were held constant across all simulations we performed. The default parameters for these simulations are included in **Table 1**.

**Table 1.** Parameters used for liquid and spatially-structured simulations displayed in **Figure 1**.

| Parameter name | Biological meaning | Default Value(s) |
| --- | --- | --- |
| growth_rate | growth rate of metabolic models | 0.125, 0.25, 0.5, 0.75, 1 |
| production_cost | cost of toxin production as a percentage of growth rate | 0.01 |
| resistance_cost | cost of resistance to toxin as a percentage of growth rate | 0 |
| toxin_coefficient | mmol of toxin produced per gram of producer biomass | 15 |
| metabolite_diff | rate of resource diffusion | 5e-6 |
| toxin_diff | rate of toxin diffusion | 5e-7 |
| space_width | simulated environment size (m x m) | 0.01 |
| spatial_seed | random seed for reproducible colony founder locations | 1, 2, 3, 4, 5, 6, 7, 8, 9, 10, 11, 12, 13, 14, 15 |
| initial_biomass | initial biomass density per colony | 2e-9 |
| grid_size | grid size for initial colony locations | 50 x 50 |
| add_signal_parameter | functional shape of how toxin alters susceptible growth rate | bounded_linear |
| conc_where_effect_ends | toxin concentration at which susceptible growth rate reaches 0 | 15 |
| max_cycles | total simulation hours | 75, unless additional time was required to reach stationary phase |
| dt | simulation time step | 5, unless a different time step was required to stabilize diffusion constants |

For spatially-structured environments, we ran either fifteen (for **Fig. 1**) or three (for **Fig. 2**) simulations of 50×50 grids, where colonies were inoculated at random locations at the start of growth. For simulations involving parameter sweeps, only three different grids were used because we found that increasing the number of replicates greatly increased computation time while generally strengthening but not changing our qualitative results. Single simulations were run for liquid experiments because, in the absence of the stochastic effect of spatial location, simulation results are deterministic. All simulations were run until biomass no longer changed between simulation steps (i.e. stationary phase was reached). Additionally, for parameter sweeps in spatially-structured environments, we ran simulations varying the mmol of toxin produced per gram of biomass, the starting frequency of susceptible colonies, the total number of founding colonies, the toxin diffusion rate, and growth rate. A full table of explored parameter space is available in **Table 2**. At the end of all simulations, we evaluated the ability of the producer to invade the total population by calculating its final population frequency. If, by the end of growth, the producer increased in frequency over its starting frequency (producer / (producer + susceptible + resistant) > 0.1, or other initial frequency as relevant), invasion was considered successful. We also evaluated the ability of the producer to outcompete resistant cheats. Because producers and cheats were seeded at a 1:1 ratio at the start of every simulation we performed (a frequency of 0.5), the producer was considered to have a higher relative fitness if its final frequency increased over 0.5 (producer / (producer + resistant) > 0.5). These two types of relative fitness values were calculated for each simulation and considered to investigate when toxin production should be maintained.

**Figure 1.**
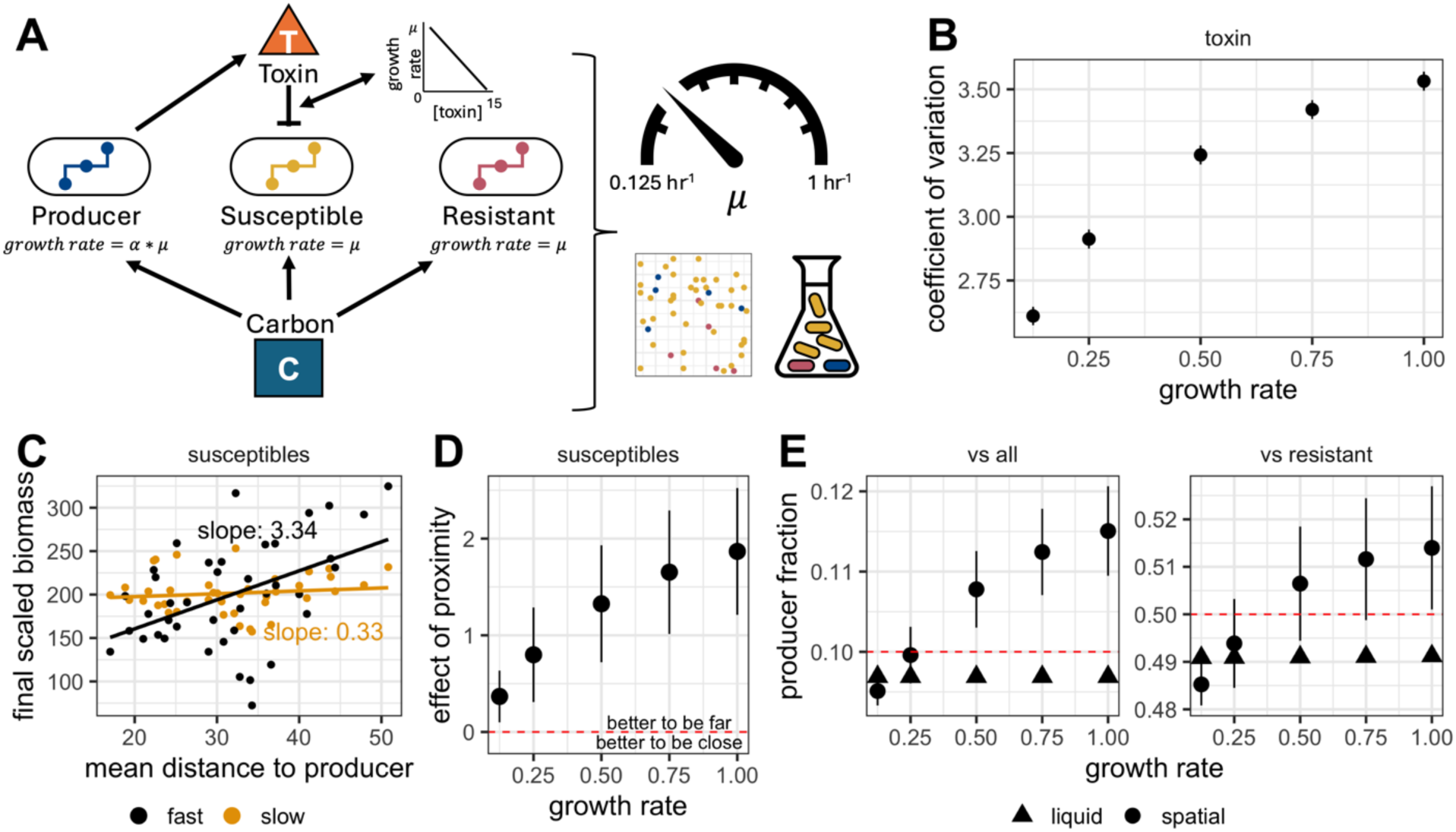
Increasing growth rate increases the localization of resource and interference competition. **A:** Schematic of simulations. Producers, susceptibles and resistant cheaters grow in a shared environment competing for a single carbon source. Producers are subjected to a cost of toxin production as a fraction of growth rate, where α represents the percentage cost and μ represents growth rate. Susceptible and resistant cheaters grow at the same rate μ. During growth, toxins are generated by producers and diffuse into the environment. Toxin reduces susceptible growth rate linearly; as toxin concentration increases to 15 mmol, susceptible growth rate is reduced to 0. Simulations are run in either a randomly generated spatially-structured environment with 50 founder colonies (grid) or a well-mixed environment (flask). All simulations are seeded with the same biomass, at a ratio of 80% susceptible to 10% resistant cheater and 10% producer. The growth rate μ is tuned between a value of 0.125 hr^-1^ and 1 hr^-1^. **B:** Maximum coefficient of variation in toxin concentration across all spatial points in a 50×50 grid over the course of growth. Points are the average of fifteen spatial replicates and error bars represent standard error of the mean. **C:** The mean distance to producer colonies for each susceptible colony plotted against final scaled colony biomass. Final biomass values at a high growth rate (1 hr^-1^) are plotted in black. Final biomass values at a low growth rate (0.125 hr^-1^) are plotted in orange. Colonies for a single representative spatial replicate are shown. **D**: The effect of proximity for susceptibles as growth rate increases. Points are the mean of fifteen spatial replicates and error bars represent standard error of the mean. Points greater than zero indicate that biomass increases with distance while points less than zero indicate that biomass decreases with distance. **E:** Producer fraction of the total population (“vs all”) or the non-susceptible population (“vs resistant”) as growth rate increases in either a liquid (triangles) or spatial (circles) environment. A producer fraction of the total population greater than 0.1 indicates successful invasion; a producer fraction of the non-susceptible population greater than 0.5 indicates higher fitness than resistant cheaters. Points are the mean of fifteen spatial replicates and error bars represent standard error of the mean. Only one simulation was run for liquid environments. *For all simulations shown: Production cost is 1% of growth, toxin coefficient (mmol toxin / gram of growth) is 15, there is no growth cost for resistance, the ratio of toxin to resource diffusion is 0.1, toxin signal parameters are bounded linear, and total environment size is 0.01 (1 cm x 1 cm). A full list of simulation parameters is available in* ***Table 1***.

**Figure 2.**
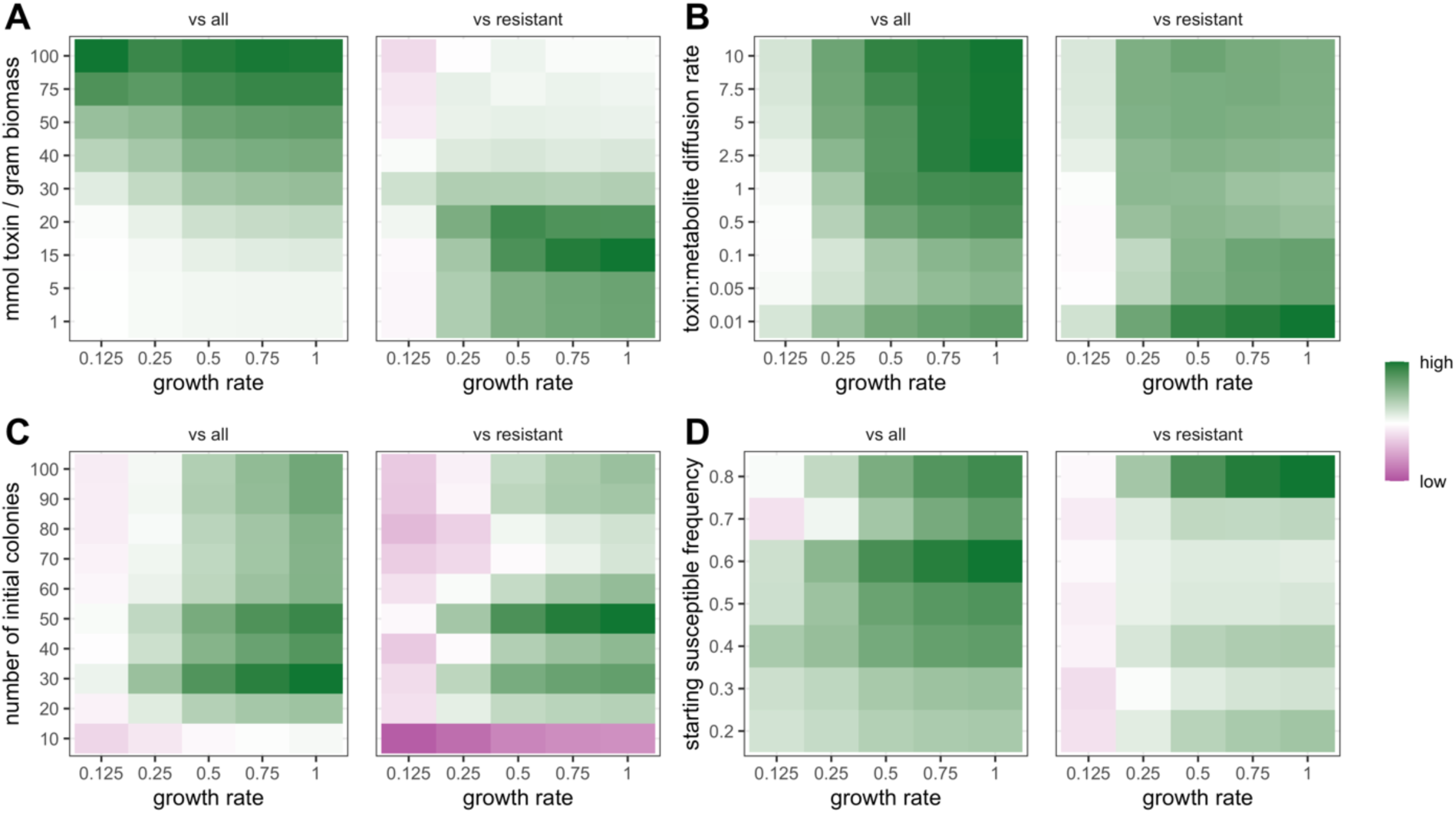
Increasing growth rate increases the strength of selection for toxin production across a wide range of parameter space. **A:** Fitness of toxin producers as growth rate and the mmol of toxin produced per gram of biomass change. **B:** Fitness of toxin producers as growth rate and the toxin diffusion rate change. The ratio of toxin diffusion rate to the rate of metabolite diffusion is plotted on the y-axis. **C:** Fitness of toxin producers as growth rate and the total number of initial colonies change. The ratio of susceptibles, producers and resistants does not vary with the number of colonies and is fixed at a percent ratio of 80:10:10. **D:** Fitness of toxin producers as growth rate and the starting frequency of susceptible colonies change. The initial frequency values used as the baseline for producer relative fitness in the total population changes with starting susceptible frequency (i.e. relative fitness threshold for the “vs all” facet when susceptible frequency is 0.8 is 0.1, relative fitness threshold when susceptible frequency is 0.6 is 0.2, etc.) *For all heatmaps: Green values indicate high relative fitness (>0.1 for vs all or other relevant value, >0.5 for vs resistant), white values correspond to no difference in relative fitness (0.1 or other relevant value, 0.5) and pink values indicate low relative fitness (<0.1 or other relevant value, <0.5). Values are the average ratio of final biomass (producer to susceptible and resistant or producer to resistant) across three spatial replicates. For all simulations shown: Production cost is 1% of growth, there is no growth cost for resistance, signal parameters are bounded linear, and total environment size is 0.01 (1cm x 1cm). A full list of simulation parameters is available in Table 2*.

**Table 2.**
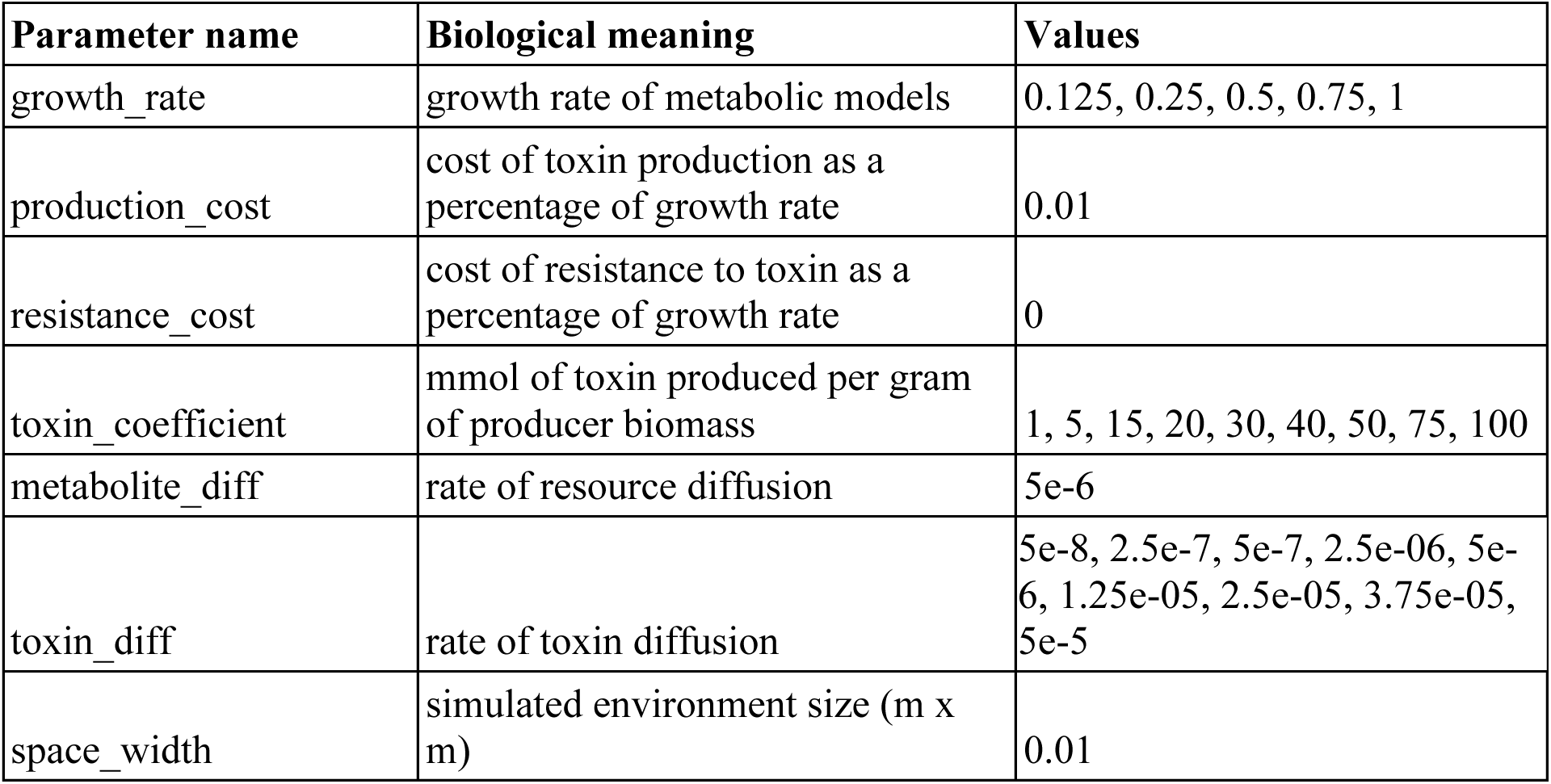
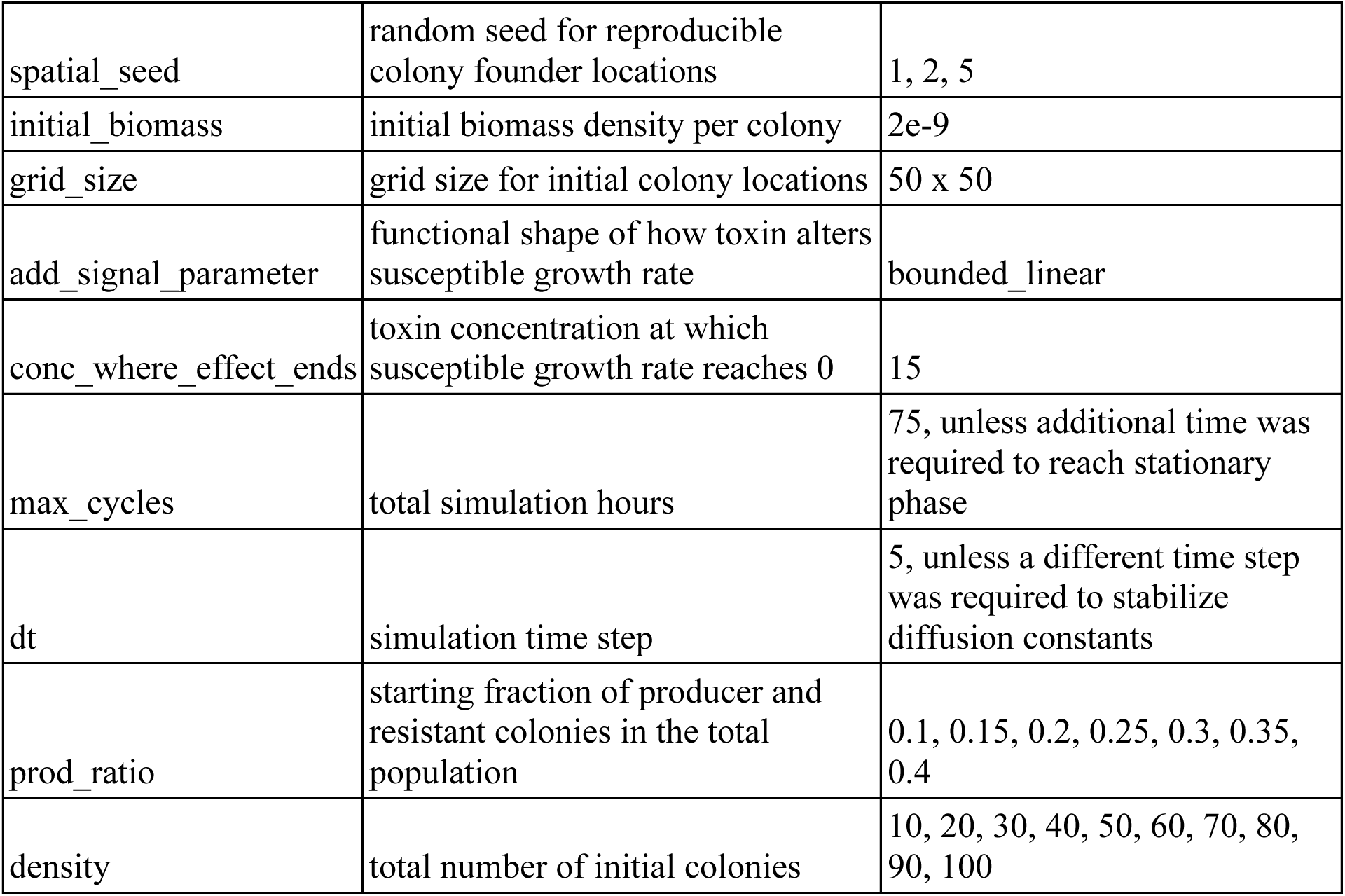
Parameter space explored for spatially-structured simulations displayed in **Figure 2**.

| Parameter name | Biological meaning | Values |
| --- | --- | --- |
| growth_rate | growth rate of metabolic models | 0.125, 0.25, 0.5, 0.75, 1 |
| production_cost | cost of toxin production as a percentage of growth rate | 0.01 |
| resistance_cost | cost of resistance to toxin as a percentage of growth rate | 0 |
| toxin_coefficient | mmol of toxin produced per gram of producer biomass | 1, 5, 15, 20, 30, 40, 50, 75, 100 |
| metabolite_diff | rate of resource diffusion | 5e-6 |
| toxin_diff | rate of toxin diffusion | 5e-8, 2.5e-7, 5e-7, 2.5e-06, 5e-6, 1.25e-05, 2.5e-05, 3.75e-05, 5e-5 |
| space_width | simulated environment size (m x m) | 0.01 |
| spatial_seed | random seed for reproducible colony founder locations | 1, 2, 5 |
| initial_biomass | initial biomass density per colony | 2e-9 |
| grid_size | grid size for initial colony locations | 50 x 50 |
| add_signal_parameter | functional shape of how toxin alters susceptible growth rate | bounded_linear |
| conc_where_effect_ends | toxin concentration at which susceptible growth rate reaches 0 | 15 |
| max_cycles | total simulation hours | 75, unless additional time was required to reach stationary phase |
| dt | simulation time step | 5, unless a different time step was required to stabilize diffusion constants |
| prod_ratio | starting fraction of producer and resistant colonies in the total population | 0.1, 0.15, 0.2, 0.25, 0.3, 0.35, 0.4 |
| density | total number of initial colonies | 10, 20, 30, 40, 50, 60, 70, 80, 90, 100 |

To quantify the localization of interactions in our spatially-structured simulations, we developed two metrics to reflect the scale of competition. First, we evaluated the maximum coefficient of variation in toxin concentration. To do so, we calculated the coefficient of variation in toxin concentration across every spatial point in our 50×50 grid for every time step up to the cessation of bacterial growth. Then, we compared the maximum coefficient of variation over the course of growth for each spatial replicate across each growth rate. We paired this with another metric of localization. For each susceptible colony, we calculated the slope of the line generated by plotting final colony biomass as a function of mean distance from producer colonies. A positive relationship between these values indicated that susceptible colonies growing further from producer colonies grew to larger sizes than those near producers, while a negative relationship indicated that susceptibles growing closer to producers tended to grow larger than those in more distant locations. We used these two values to consider how various other parameters interacted with growth rate to tune the scale of interactions.

### Curation of bioinformatics datasets

To investigate the relationship between toxin production and growth rate in natural communities, we curated two bioinformatics datasets from separate sources. First, we downloaded all the relevant genomes from the 2010 Vieira-Silva and Rocha paper “The Systemic Imprint of Growth and Its Uses in Ecological (Meta)Genomics” for which maximum doubling times were measured experimentally [13]. We used the ncbi_datasets command line tool (ncbi_datasets v. 16.24.1) to download the reference genomes (--reference, both fna and gbff files) using the strain name provided in the paper [15]. For species for which a reference genome was not found, in most cases due to changes or updates to genus names, we manually searched for the relevant accession numbers by searching NCBI’s assembly database in August 2024. If more than one complete genome was available for a given strain, we chose the first one. Of the 215 genomes experimentally validated in Vieira-Silva and Rocha, we were able to download 191 complete reference genomes from NCBI. These genome gbff files were then used as input for antiSmash v. 6.1.1 with “relaxed settings” [16]. If genome files were not annotated, gene predictions were generated using prodigal. antiSmash html outputs were imported into R v. 4.1.2 and cleaned. GenBank fna files were used as input for gRodon v. 2.0.0, which predicts maximum doubling times for prokaryotes and microbial eukaryotes based on codon usage bias and a model trained using a database of experimentally-verified doubling times [17]. Outputs were aggregated into a single dataset that included genome name, gRodon-predicted doubling time, experimentally-determined doubling time, and all relevant antiSmash-predicted biosynthetic gene clusters.

While the previous dataset included experimentally-verified growth rates, we expanded our dataset to include pure bioinformatic predictions on a larger set of 2,912 *Streptomyces* complete genomes from GenBank (both fna and gbff files) using the ncbi_datasets command line tool (ncbi_datasets v. 16.24.1) in August 2024 [15]. GenBank fna files were used as input into gRodon v. 2.0.0 [17]. GenBank gbff files were used as input into antiSmash v. 6.1.1 [16] with default settings, including “relaxed” thresholds for identifying biosynthetic genome clusters. If genome files were not annotated, gene predictions were generated using prodigal. antiSmash html outputs were imported into R v. 4.1.2 and cleaned. Outputs were aggregated into a single dataset that included genome accession number, gRodon-predicted doubling time, and all relevant antiSmash-predicted secondary metabolite clusters. For both datasets, a full list of taxon names or accession numbers used for downloads from NCBI is available in **Supplemental Table 1.**

### Bioinformatics dataset analysis

After curating our two datasets, we statistically evaluated the relationship between growth rate (independent variable) and biosynthetic gene cluster presence (dependent variable) in R v. 4.1.2. Growth rates in hr^-1^ were calculated based on doubling times, such that maximum growth rate = ln(2) / (maximum doubling time in hours). To assess whether growth rates were associated with the presence of a biosynthetic gene cluster in a genome, we fit a generalized linear model to each cluster type using the argument “binomial” and link “logit”, which is appropriate for predicting the probability of an occurrence (i.e. the probability that a biosynthetic gene cluster is present as a function of growth rate). Experimentally-determined doubling times were used to calculate growth rates for model generation for the Vieira-Silva and Rocha dataset, while gRodon-predicted doubling times were used for growth rates for the *Streptomyces* dataset. A separate test was applied to each biosynthetic gene cluster. We corrected for multiple comparisons across biosynthetic gene clusters using a Benjamini-Hochberg correction. 95% confidence intervals were calculated for β coefficients using the R package MASS v. 7.3.

Because there is uncertainty in the growth rate predicted by gRodon (**Supplemental Fig. 1**), we evaluated the robustness of our *Streptomyces* results using two resampling approaches (**Supplemental Fig. 2**). First, for each genome, we randomly sampled 10,000 new doubling times from a uniform distribution using the 95% confidence interval provided by gRodon for each prediction to set the minimum and maximum of the distribution (minimum = lower CI, maximum = upper CI). For our second approach, we randomly sampled 10,000 new doubling times from a normal distribution, where the mean of the distribution was the original prediction provided by gRodon and the standard deviation was calculated from the 95% confidence interval (mean = gRodon-predicted doubling time, sd = (sqrt(sample size)*(upper CI – lower CI)) / (2*1.96)). Following resampling, we generated logistic regression models between biosynthetic gene cluster presence and growth rate and evaluated their significance as previously described.

We compared the number of statistically significant models for each resampling approach, the proportion of total bootstrapping replicates in which the presence of a given biosynthetic gene cluster was significantly correlated with growth rate, and the average β coefficient across all bootstrapping replicates to our initial results (**Supplemental Fig. 2**). BGC classes which were significantly correlated with growth rate in at least 95% of bootstrapping replicates for both resampling approaches were considered robust correlations (**Fig. 5B**). We also evaluated the robustness of our comparative bioinformatics approaches for the Vieira-Silva and Rocha dataset by rerunning the previously described analyses using gRodon-predicted doubling time (**Supplemental Fig. 3**) or without genomes with no predicted BGCs (**Supplemental Fig. 4**).

## RESULTS

### Increasing growth rate reduces the scale of resource and interference competition

We tested the effect of growth rate on fitness of a toxin producer by conducting simulations in either spatially-structured or well-mixed environments using the COMETS platform (**Fig. 1A**) while varying the entire community’s growth rate (μ) between 0.125 hr^1^ and 1 hr^-1^. Producer models paid a modest growth cost for production (α = 0.01 or 1% of growth).

Fifteen spatial replicates were performed, with varying initial biomass locations. We tracked the biomass of individual founder colonies as well as the concentrations of toxin and carbon over space and time. The rest of the default parameters used for these simulations are available in **Table 1** and all other modeling constraints are described in the **Methods**.

We first found that increasing growth rate increased spatial heterogeneity in toxin concentration. We calculated the coefficient of variation in toxin concentration across the entire environment for each time step during growth at different rates and identified the maximum coefficient for each growth rate across all time steps. In doing so, we observed that this value increased with growth rate (**Fig. 1B**), in part because at high growth rates, some individual locations experienced local toxin concentrations higher than those experienced at any location when growth rate was low. This suggested that higher growth rates increased interaction localization by driving production of higher concentrations of toxin in individual time steps and reducing the rate of toxin dilution relative to growth.

We next tested how growth rate influenced the fitness of a susceptible colony as a function of its proximity to the toxin producers. Specifically, we calculated the mean distance to producer colonies for each susceptible founder and considered how that distance impacted final biomass of the susceptible colony. (**Fig. 1C**). Mean distance was chosen to capture the cumulative effects of toxin exposure from all toxin producers present in the environment. Using this metric, we found that, at a low growth rate, there was no relationship between proximity to producer colonies and susceptible colony biomass (**Fig. 1C**). In contrast, at a high growth rate (**Fig. 1C**), being farther from producer colonies was correlated with an increase in growth (**Fig. 1C**). We called the slope of this relationship the effect of proximity and calculated it for all fifteen replicate spatial arrangements across all growth rates (**Fig. 1D**). We found that, as growth rate increased and the length scale of interactions decreased, susceptibles paid a greater price for their initial proximity to producers. Together, the coefficient of variation in toxin concentration and the effect of proximity on susceptible biomass underscored that growth rate changed the interaction scale of competition, in alignment with our initial hypotheses.

Having found that the fitness of susceptible colonies varied with growth rate due to changes in the length scale of interactions, we next explored how varying growth rate influenced a producer’s overall fitness and its fitness relative to a resistant cheater genotype. To do so, we compared the final biomass and species ratios of our spatial simulations with simulations in liquid. Simulations in well-mixed environments were performed only once because results were deterministic (i.e. independent of starting spatial location). We calculated both the fraction of the final population biomass composed of the producer relative to the starting frequency of 10%, as well as the relative fitness of the producer against the resistant cheater. Relative fitness was measured as the final frequency of the toxin producer relative to the resistant (producer / (producer + resistant)) and compared to the initial frequency of 0.5. Therefore, any final frequency above 0.5 indicated selection for toxin production. We found that in a liquid environment, toxin production was not selected for by either of our metrics, regardless of growth rate (**Fig. 1E**). In contrast, in a spatially-structured environment, we observed a positive relationship between growth rate and the fitness of the producer population (**Fig. 1E**). At low growth rates (0.125 - 0.25 hr^-1^), producers were not able to invade the total population nor outcompete resistant cheaters (**Fig. 1E**). The relative fitness of toxin production increased with maximum growth rate, with production selected for using both metrics at growth rates of 0.5 hr^-1^ and above (**Fig. 1E**). Our results therefore aligned with our hypothesis that spatial structure should be more likely to maintain toxin production at higher growth rates.

### Growth rate is consequential for selection on toxin production over a range of parameter space

To test the robustness of our results, we ran simulations varying four additional parameters concurrently with growth rate. Specifically, we varied the mmol of toxin produced per gram of biomass (1 to 100 mmol / gram), the rate of toxin diffusion relative to the rate of metabolite diffusion (0.01x to 10x the rate of metabolite diffusion), the initial fraction of the population composed of susceptible colonies (20 to 80% of the total population), and the total number of initial colonies (10 to 100 total colonies). The toxin concentration, toxin diffusion rate, and initial population density were manipulated to examine different variables contributing to the length scale of interactions. The starting frequency of susceptibles was manipulated to consider how changing the likely identity of nearest neighbors would impact the benefit of increasing growth rate. For all simulations, we again varied the community’s growth rate (μ) between 0.125 hr^1^ and 1 hr^-1^ and kept the cost of production at 1% of growth. Three randomly initialized spatial layouts were completed for each condition. In our simulations varying the initial susceptible fraction, producers and resistant cheaters were always seeded at the same 1:1 ratio, but they represented increasing fractions of the total population, ranging from 10% of the total initial population each (5 resistant and 5 producer colonies) to 40% each (20 resistant and 20 producer colonies). A full list of simulation parameters is available in **Table 2**.

We found that growth rate increased the strength of selection for toxin production across a wide parameter space. We found that higher growth rate generally increased the benefit of toxin production along both our metrics of fitness even as other parameters determined the direction and magnitude of selection (**Fig. 2A-D**). Consistent with previous work [9], we found that toxin production provided the biggest benefit at intermediate levels of toxin production and colony density (**Fig. 2A, 2C**). However, our results demonstrated that the strength of this effect was strongly influenced by growth rate. Our results underscored that growth rate is just one of many potential factors influencing selection on toxin production, but that in nearly all cases, even as other biophysical and environmental parameters vary, higher growth rates should increase the benefit of toxin production relative to slower growth by increasing interaction localization.

### Growth rate correlates with the presence of certain biosynthetic gene clusters in diverse datasets

To understand whether, as our modeling predicted, faster microbial growth rates were more likely to select for toxin production in natural communities, we curated two datasets of genomes from different sources. There are measurable genomic signatures of many species’ maximum growth rate, such as the number of *rrn* gene copies and the codon usage bias of these genes [13, 17–19]. The software gRodon uses these genomic signatures in combination with a model trained based on a database of experimentally-verified doubling times to generate predictions of a species’ growth rate, which we leveraged for our analysis [13]. Likewise, there are genomic signatures of toxin production. Because they are not strictly necessary for growth, toxins are classified as secondary metabolites and are often produced by groups of enzyme-encoding genes that cluster together within a genome, known as biosynthetic gene clusters (BGCs) [20]. These clusters can be detected using existing software tools such as antiSmash [16]. We therefore used genomic predictions of both maximum growth rate and the presence of various BGCs to investigate the relationship between toxin production and growth rate in a diversity of strains.

Our datasets were derived from the growth rate database from Vieira-Silva and Rocha (2010) [13] and nearly 3,000 complete *Streptomyces* assemblies available from NCBI GenBank. The dataset from Vieira-Silva and Rocha (**Fig. 3**) spanned 191 genomes. Those genomes were predicted by antiSmash to contain between 0 and 50 BGCs across 49 cluster types, with a mean of 4.3 clusters per genome and a median of 2 (**Fig. 3A**). All genomes were experimentally-determined to have a maximum growth rate (ln(2) / observed doubling time, in hr^-1^) between 0.003 and 3.47, with a mean of 0.51 and a median of 0.21 (**Fig. 3B**; Vieira-Silva and Rocha 2010). We also curated a dataset of 2,912 *Streptomyces* genomes available on NCBI. All genomes included in this dataset were predicted by antiSmash to contain between 3 and 164 BGCs, representing 56 different types of clusters, with a mean of 47 BGCs per genome and a median of 45 (**Fig. 3A**). All *Streptomyces* genomes had gRodon-predicted growth rates (ln(2) / predicted doubling time, in hr^-1^) between 0.15 and 1.26, with a mean of 0.61 and a median of 0.60 (**Fig. 3B**). Additionally, certain types of BGCs were highly represented across all our tested genomes. For example, 40% of genomes in the Vieira-Silva and Rocha dataset and 100% of *Streptomyces* genomes were predicted by antiSmash to contain at least one terpene BGC (**Fig. 3C**). RiPP-like, siderophore, and NRPS BGCs were also predicted to be present in a significant percentage of genomes in each dataset (**Fig. 3C**).

**Figure 3.**
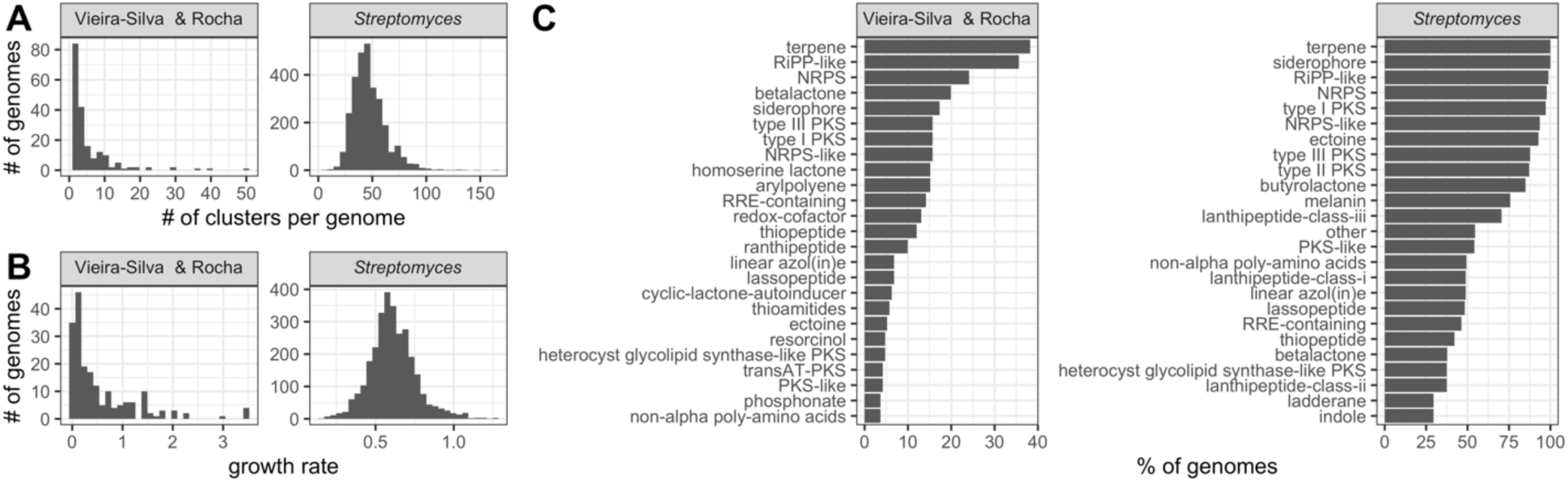
Two genomics datasets drawn from different sources cover a great deal of microbial diversity in terms of number of biosynthetic gene clusters per genome and growth rates. **A:** Histogram of total number of predicted biosynthetic gene clusters per genome for each dataset. **B:** Histogram of maximum growth rates calculated from doubling times for each dataset (observed for Vieira-Silva and Rocha, predicted for *Streptomyces*). **C:** Percent of genomes containing a given type of biosynthetic gene cluster per dataset. Only the 25 most represented types are displayed for each dataset, respectively.

We used our datasets to test the relationship between the presence of secondary metabolite gene clusters and growth rate. We generated generalized logistic regression models (binomial family, logit link) and used Benjamini-Hochberg p-value adjustment for multiple comparisons. Despite the relatively lower power of the analysis (due to 49 statistical tests [one per BCG] and 191 datapoints [genomes]) and potential confounding variables including phylogenetic distance, we found that the presence of five types of BGCs were significantly related to growth rate in the Vieira-Silva and Rocha (2010) [13] dataset (**Fig. 4A**). The probability of the presence of all five of these types of biosynthetic gene clusters - siderophores, ectoines, betalactones, nonribosomal peptide synthetase clusters (NRPS), and lanthipeptide class I clusters - were predicted to increase with growth rate (**Fig. 4B-C**), such that they had a higher probability of appearing in faster-growing strains. Of these BGCs, betalactones, non-ribosomal peptide synthetase clusters, and lanthipeptide class I BGCs can all encode for natural products with antimicrobial activity (**Fig. 4B**) [21–23].

**Figure 4.**
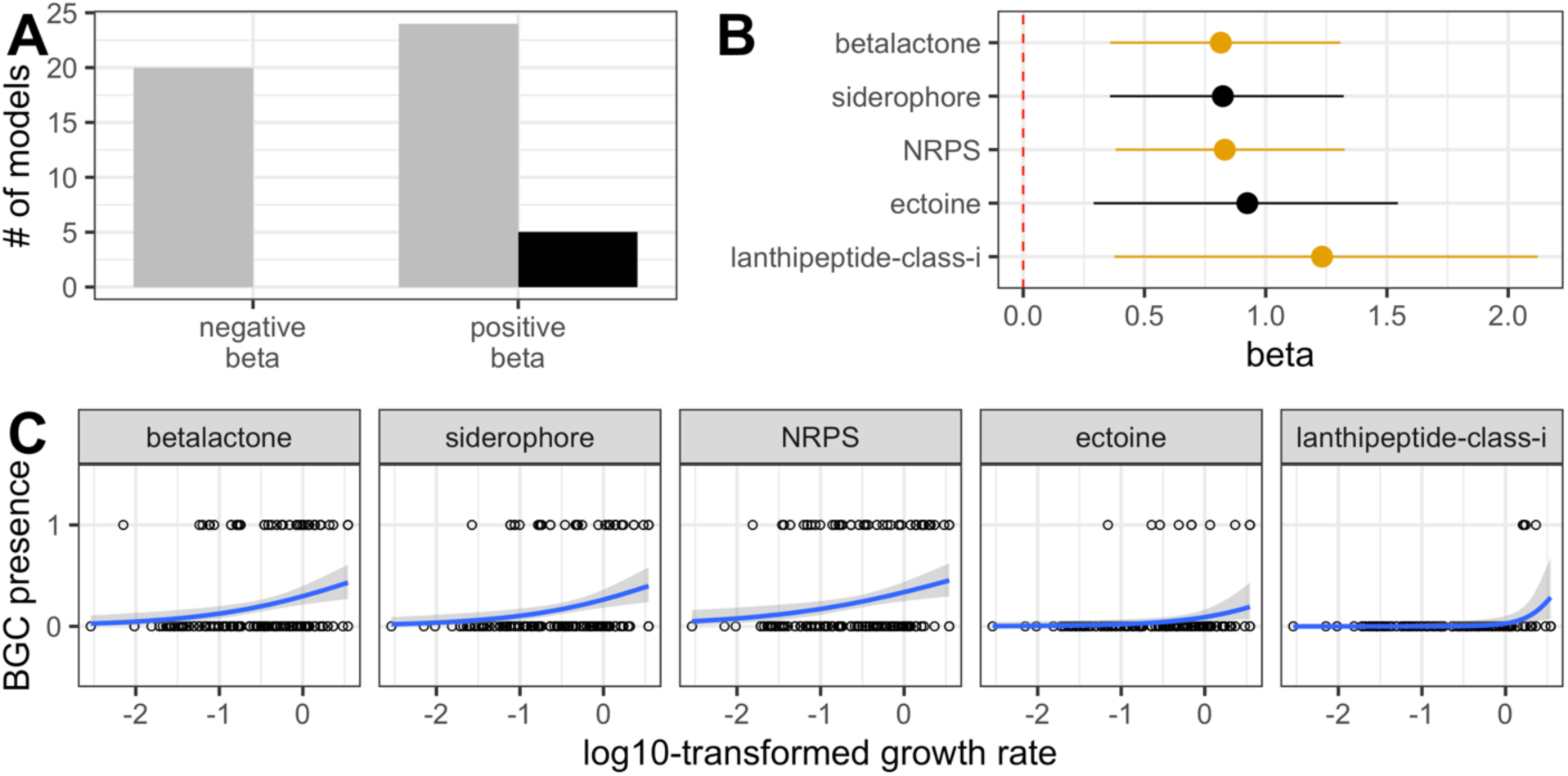
In the Vieira-Silva and Rocha dataset, the presence of five types of biosynthetic gene clusters is more likely as growth rate increases. **A:** Number of logistic regression models with negative and positive β values. The number of models with significant correlations (following Benjamini-Hochberg correction) are colored in black. **B:** β coefficients for logistic regression models generated for growth rate and BGC class presence with significant adjusted p-values. Line segments indicate 95% confidence intervals for model estimates. BGC classes that can potentially encode antimicrobial compounds are colored in orange. **C:** Regression plots of the five BGC classes with a significant correlation between presence and growth rate, in order of increasing β coefficient. Growth rates are log10-transformed to better observe the range of small values because growth rates across the dataset are left-skewed.

Our simulation results were also consistent with comparative genomic analyses of our *Streptomyces* dataset, where we found that the presence of 29 secondary metabolite gene types was significantly correlated with growth rate (**Fig. 5A**). The probability of the presence of six of these BGC classes increased with decreasing growth rate (β < 0), while the probability of the presence of 23 BGC classes were predicted to increase with increasing growth rate (β > 0), (**Fig. 5B-D**). After evaluating the robustness of our statistical models to account for the predictive error of gRodon, we found that 19 classes of BGCs were significantly correlated with growth rate in at least 95% of 10,000 bootstrapping replicates following two resampling approaches, indicating the robustness of the relationship: homoserine lactone, RRE-containing, arylpolyene, indole, thiopeptide, other, linear azol(in)es, transAT-PKS-like, polyketide synthase (PKS) type I and II, non-alpha poly-amino acids, aminoglycoside, lanthipeptide class IV, butyrolactone, NRPS, ranthipeptide, phosphoglycolipid, melanin, and RiPP-like (**Fig. 5B**). For all 19 BGC classes, the direction of the correlation was robust to resampling (β < 0 or > 0), with four classes negatively correlated with growth rate and 15 positively correlated. Of the 15 types of BGC classes positively correlated with growth rate, 10 can or generally do encode natural products with antimicrobial properties: RiPP-like, NRPS, butyrolactone, lanthipeptide class IV, PKS types I and II, aminoglycoside, thiopeptide, transAT-PKS-like, and linear azol(in)e-containing peptides clusters (**Fig. 5B**) [21–32]. Of the four BGC classes negatively correlated with growth rate, indoles and RRE-containing (RiPP recognition element) clusters can encode secondary metabolites that might act as toxins (**Fig. 5B**) [33–35].

**Figure 5.**
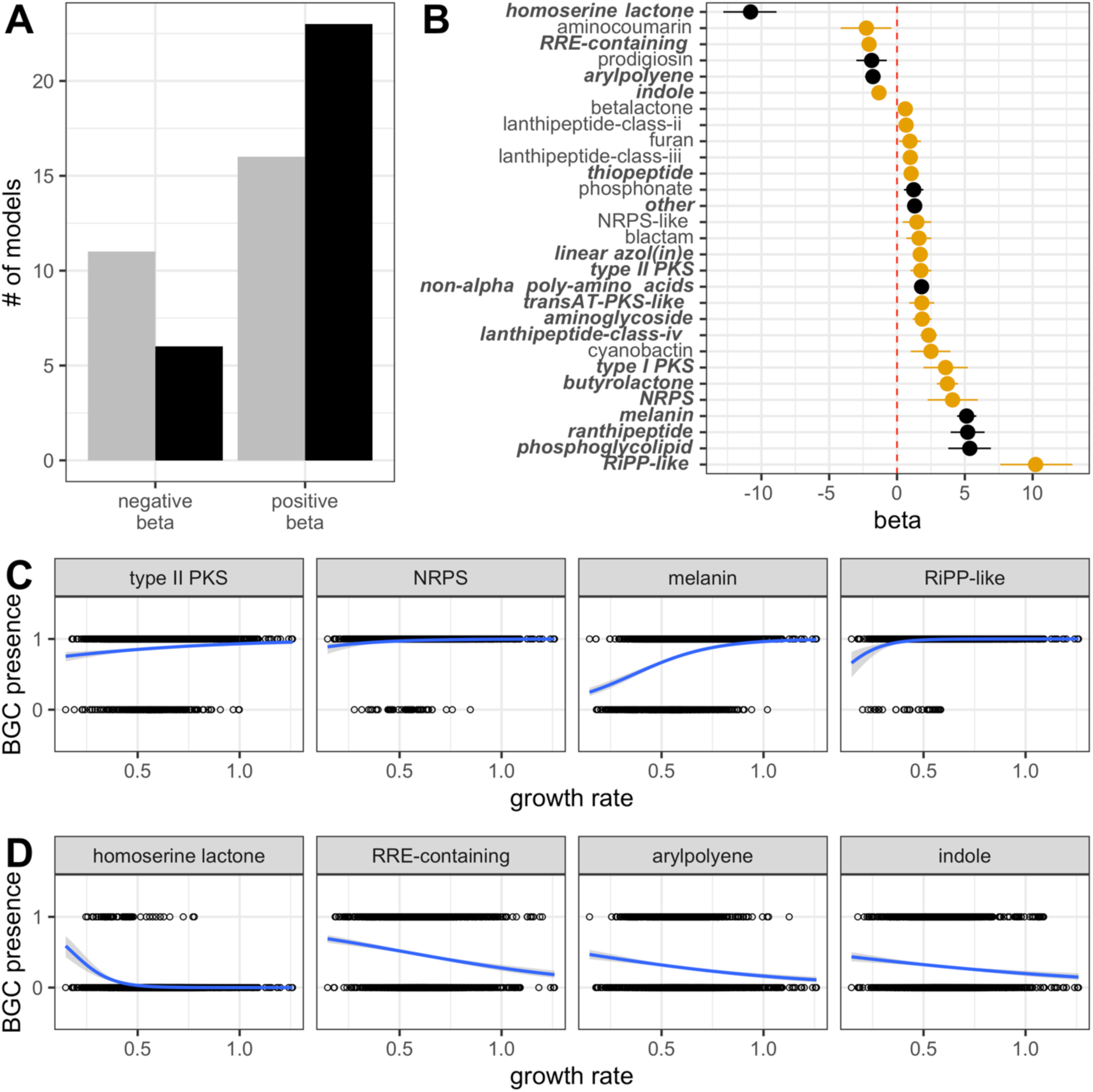
In the *Streptomyces* dataset, growth rate robustly correlates with the presence of 19 BGC classes. **A:** Number of logistic regression models with negative and positive β values. The number of models with significant correlations (following Benjamini-Hochberg correction) are colored in black. **B:** β coefficients for logistic regression models generated for growth rate and BGC class presence with significant adjusted p-values. Line segments indicate 95% confidence intervals for model estimates. BGC classes that can potentially encode antimicrobial compounds are colored in orange. The names of BGC classes with correlations that are robust following resampling are bolded and italicized on the y-axis. **C:** Regression plots of four representative BGC classes with a robust significant positive β coefficient, in order of increasing β coefficient. **D:** Regression plots of all four BGC classes with a significant negative β coefficient that is robust to resampling, in order of increasing β coefficient. *For all data shown:* β *estimates and p-values displayed were calculated using the original predicted doubling time value provided by gRodon, without resampling*.

## DISCUSSION

Our study integrated a genome-scale metabolic modeling framework with bioinformatic analyses to investigate the relationship between growth rate and selection for toxin production. Consistent with previous work [2, 6–7], our simulations showed that spatial structure is required to select for toxin producers when producers invade from rare. Our work builds upon existing theory by demonstrating that increasing microbial growth rates increases the strength of selection for toxin production by increasing the localization of interactions. Our simulations also showed that increasing growth rate increased the benefit of toxin production even as other biophysical and environmental parameters tuned the direction and strength of selection. Consistent with our simulations, our comparative genomics approaches revealed correlations between growth rate and the presence of multiple types of biosynthetic gene clusters across two microbial datasets.

Together, our results suggested that toxin production should be more beneficial and therefore more prevalent in spatially-structured environments where microbes are growing rapidly.

Using COMETS simulations, we showed that faster growth increased the benefit of toxin production because faster growth increased environmental heterogeneity and interaction localization. Specifically, increased heterogeneity at higher growth rates increased the benefit of production against susceptible strains by reducing toxin dilution, ensuring that higher concentrations reached susceptible colonies within a producer’s neighborhood. In comparison, at low growth rates, susceptible colonies experienced relatively low and homogenous concentrations of toxin across the environment. While our simulations were performed based on an assumption of a two-dimensional environment, evidence exists that increased toxin localization is even more critical for the success of toxin producers in three-dimensional space [36]. Additionally, increased interaction localization increased the benefit of toxin production relative to resistant cheaters because the benefit of toxin production remained spatially restricted to the producer. At high growth rates, inhibition of nearby susceptibles allowed toxin producers to make more effective use of local resources because these nutrients were unlikely to diffuse to distant colonies before stationary phase was reached. Resistant cheaters could only make use of these resources if they were within the interaction neighborhood of the producer, which became less likely as increasing growth rate shrank neighborhood size. This result aligned with theory on the evolution of public goods production [37], suggesting that in spatially-structured microbial communities, high growth rate can help offset the tragedy of the commons for toxin producers.

There are several caveats in our modeling approach. First, we chose to model toxins as bacteriostatic rather than bactericidal [38]. This distinction may not significantly influence the abundance of resources available to producers, though this would depend on the mechanism of action for the toxin. For example, if bactericidal toxins lyse cells during killing, cellular debris could serve as an additional resource for surviving strains, increasing resource availability [39]. On the other hand, if bactericidal toxins result in death but not lysis, the resources that are assimilated by susceptibles prior to death would not be available to producers, and results might broadly align with our findings using bacteriostatic toxins. That said, bactericidal toxins may generally be more consequential for the overall strength of selection for toxin production by significantly changing the frequency of susceptibles. We also chose to model constitutive toxin production, while many microbial strains only produce antimicrobial compounds in response to environmental signals [40]. Differential production may ultimately reduce production-associated costs and limit the benefits to resistant cheaters, which could strengthen selection for toxin production. However, we anticipate that the relationship between high growth rate and increased benefit of toxin production should largely hold even when toxins are not produced constitutively. Finally, our results focus solely on the benefits of toxin production within a single period of growth. However, there are some bacterial strains, *Streptomyces* in particular, that primarily produce toxic secondary metabolites when transitioning to sporulation [41]. Our approaches are not designed to investigate hypotheses around a “future defense” strategy, where toxins do not provide a significant benefit in a single period of growth but their impact instead becomes important over a series of resource fluctuations. In such a case, it is possible low growth rates would select more strongly for toxin-producing classes of BGCs. This is because when growth rates are slow, interference competition is less effective, as we have shown. However, in such a scenario, secreting toxins at the end of a growth phase could ensure better access to nutrients in the future when new resources become available by preventing the resumption of growth by susceptible microbes. Our bioinformatics approach suggests that this may be relevant for the presence of some types of BGCs, as we did find two classes of secondary metabolite clusters that can encode compounds with antimicrobial activity that were represented more frequently in slow-growing genomes [33–35]. Future studies could evaluate such a hypothesis by extending our model into a framework that considers temporal fluctuations in resource availability.

In alignment with our modeling results, our bioinformatic analyses suggested a correlation between maximum microbial growth rate and the presence of multiple biosynthetic gene clusters. These correlations emerged even though this type of analysis should be subject to a great deal of noise present both in the prediction of maximum growth rates (**Supplemental Fig. 1-3**) and BGC identification (**Supplemental Fig. 4-5**). Additionally, the datasets that we used for this approach include bacteria from diverse environments and genera that will differ along many axes other than growth rate. The ability of these analyses to identify any significant correlation at all is therefore remarkable and broadly supportive of our simulation results. While not all biosynthetic gene clusters of any given type will encode toxins, many of the cluster types that we found to correlate positively with growth rate can in fact be involved in interference competition. For example, some betalactone [21], NRPS [22] and lanthipeptide class I [23] BGCs can produce compounds with potent antimicrobial activity. The fact that fast-growing genomes were more likely to harbor these types of BGCs was supportive of our hypothesis that increasing growth rate should select more strongly for toxin production. However, some BGCs that correlated positively with growth rate are not typically associated with toxin production. In the Vieira-Silva and Rocha dataset, both siderophore and ectoine clusters were more likely to appear in fast-growing strains, though neither are associated with antimicrobial activity [42–43]. This demonstrates that there were other variables we did not explore that may correlate with BGC presence. Several rapidly-growing *Vibrio* strains - common in estuarine communities where ectoines can improve fitness under high osmolarity conditions [44] - were predicted to harbor ectoine BGCs in the Vieira-Silva and Rocha dataset. In this case, the positive correlation between BGC presence and growth rate we observed is likely a result of environmental conditions. Our comparative genomics approach also underscored that some types of BGCs may be more likely to be present in slow-growing strains. In our *Streptomyces* dataset, BGCs encoding quorum-sensing molecules called homoserine lactones [45] occurred more frequently in slow-growing strains, a finding that aligns with previous work in *Pseudomonas aeruginosa* demonstrating that acyl homoserine lactones accumulate rapidly when growth rate is low [46].

Finally, many of the BGCs in our datasets did not correlate with growth rate at all, regardless of whether they can encode toxins. This is unsurprising given that genomes are subject to myriad selection pressures and drift that accumulate for generations; there may be many reasons a genome that would theoretically benefit from toxin production does not harbor any toxin-encoding BGCs. We therefore expect that our bioinformatics results are supportive of the positive relationship between growth rate and selection for toxin production, but not definitive.

Our results lay the foundation for multiple routes of fruitful research. First, our findings suggest that the hunt for natural products should focus on spatially-structured environments where strains are likely to grow rapidly. This stands somewhat in contrast to the vast number of planktonic marine microbial species that harbor toxins and other natural products [47], although we note that marine snow, organic matter that serves as a spatially-structured, relatively resource-rich environment for associated bacteria as it falls toward the ocean floor [48], might be a particularly good place to search for toxin producers. Additionally, we expect that many other variables beyond growth rate will impact selection for toxin production. For example, our work has focused on antimicrobial compounds as toxins, while there are scenarios under which they are better understood as signaling molecules. Though many natural products have direct inhibitory effects, some organisms instead respond to these compounds by modifying their growth strategies or producing their own natural products [49]. More complicated community dynamics are likely to emerge in cases where secondary metabolites are signals for some subset of a community and inhibitory compounds for others, and the selective pressure to maintain toxin-producing biosynthetic gene clusters may vary as a result. Furthermore, as we have shown, in spatially-structured environments, the extent of interaction localization - or how many neighbors a microbe experiences – will depend on many variables [11]. In natural environments such as the soil, the connectivity between distant neighborhoods can be influenced by environmental properties like rainfall and soil grain size [50–51]. Yet it is not clear how these properties change with environment type, or even within microenvironments in a single area. We suggest that additional studies focused on the biophysical properties of spatially-structured environments will improve our ability to predict the conditions under which toxin production should be maintained, which will increase our ability to predict and engineer microbial communities and harness natural products. To aid in these studies, we have now generated an open-source tool that will help facilitate greater investigation of the critical link between microbial metabolism and selection for secondary metabolites such as toxins.

## Supporting information

Supplemental Material Descriptions

## ACKNOWLEDGMENTS

The authors thank E. Takano, J. Connelly, R. Breitling, F. del Carratore, and L. Kinkel for thoughtful discussions that led to the conception of this manuscript. They also thank S. Hammarlund, L. B. Smith, J. N. V. Martinson, X. Xiong, H. Ahmed and A. Lee for helpful comments throughout the execution of the project. This work was supported by NSF 1935458 to WRH and MJS.

## DATA AVAILABILITY STATEMENT

Data analysis, statistics, and figure generation were performed in R v. 4.1.2 using custom scripts available at https://github.com/bisesi/Toxins-Growth-Rate. Shell scripts to download taxonomic data and run command line antiSmash analyses, along with Python scripts to run COMETS simulations, are available at the same link. A vignette outlining use of the COMETS toxin/signaling extension is available at https://bisesi.github.io/Toxins-Growth-Rate/.

