## Supplemental Material Descriptions for "The strength of selection for costly toxin production increases with growth rate"

**Supplemental Table 1.** Csv file containing the full list of taxa names or species IDs downloaded from NCBI using the command line tool ncbi_datasets for each dataset. Accession numbers were used to download *Streptomyces* genomes. Taxon names were used to download genomes for the Vieira-Silva and Rocha and the Toxinome datasets.

**Supplemental File 1.** PDF file containing four supplemental figures (**Supplemental Fig. 1-4**), appropriate legends and captions, and additional analysis information related to the predictive ability of the package gRodon and the robustness of our bioinformatics results.

**Supplemental File 2.** PDF file containing one supplemental figure (**Supplemental Fig. 5**), the corresponding legend and captions, additional analysis information, and references for analysis of the relationship between toxin gene presence and growth rate in the Toxinome database.
