## Supplemental File 1 for "The strength of selection for costly toxin production increases with growth rate"

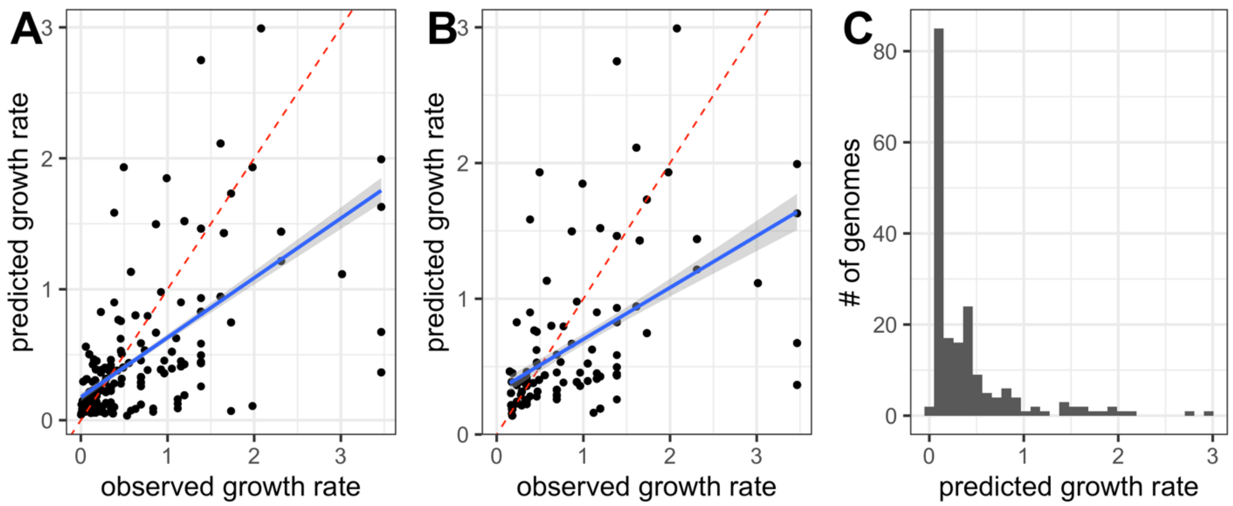


**Supplemental Figure 1. The Vieira-Silva and Rocha dataset demonstrates the predictive limitation of gRodon. A:** Scatterplot of gRodon predictions of growth rate versus observed growth rates for the Vieira-Silva and Rocha dataset. The dashed red line represents perfect predictions, while the blue line corresponds to the linear regression between values, with an estimate of 0.455 and R-squared of 0.425. While gRodon is reported to underestimate actual maximum doubling times, with predictions being most accurate when doubling times are less than 5 hours (a growth rate of 0.14 $hr^{-1}$), this analysis suggested that that gRodon in fact generally overestimated doubling time, therefore underestimating growth rate**. B:** Scatterplot of predicted growth rate versus observed growth rates for genomes with a doubling time of less than 5 hours (growth rate > 0.14 $hr^{-1}$) for Vieira-Silva and Rocha dataset. The dashed red line represents perfect predictions, while the blue line corresponds to the linear regression between values, with an estimate of 0.38 and an R-squared of 0.26. Removing doubling times greater than 5 hours (i.e. growth rates less than 0.14 $hr^{-1}$) reduced the fit between predicted and observed growth rates. **C:** Histogram of gRodon-predicted growth rates for Vieira-Silva and Rocha dataset (mean 0.41, median 0.21, min 0.035, max 3.0). Predicted growth rates did not capture either the maximum or minimum values of observed growth rates. We therefore expected that the use of gRodon-predicted rates for the *Streptomyces* dataset should somewhat overestimate the probability of the presence or absence of a given biosynthetic gene cluster by truncating the true variance in growth rate**.**

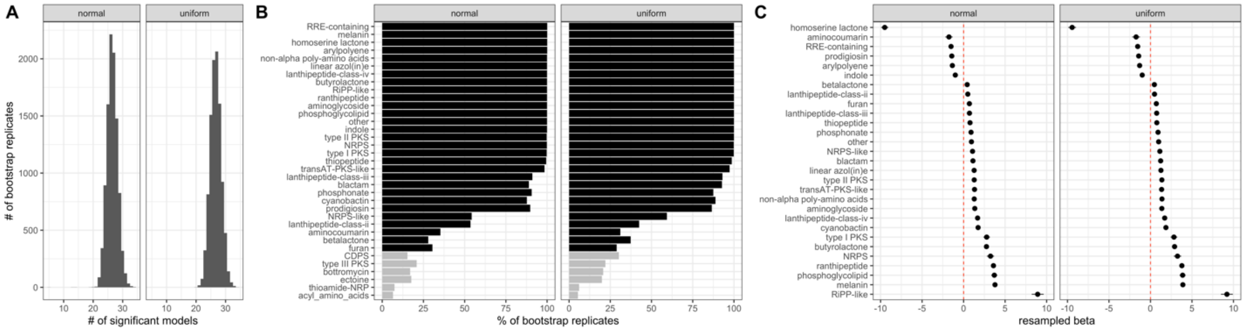


**Supplemental Figure 2. In the *Streptomyces* dataset, 19 classes of BGCs are robustly correlated with growth rate following two different bootstrapping approaches. A:** Histogram of total number of statistically significant models across 10,000 bootstrapping replicates for two resampling approaches. Using a uniform distribution for resampling, the median number of significant models per replicate is 27, with a mean of 26.7, a maximum of 34, and a minimum of 20. Using a normal distribution for resampling, the median number of significant models per replicate is 26, with a mean of 26.5, a maximum of 34, and a minimum of 5. **B:** Percent of times the presence of a BGC class is significantly correlated with growth rate across 10,000 bootstrapping replicates for two resampling approaches. Only BGC classes that were significant in >5% of bootstrapping replicates for at least one resampling approach are shown. BGC classes that were significantly correlated with growth rate using our initial dataset without resampling are colored in black (**Fig. 5B**), and classes that were not significantly correlated with growth rate in our initial dataset without resampling are colored in gray. The 19 classes of BGCs that are significantly correlated with growth rate in at least 95% of bootstrapping replicates for both resampling approaches are considered significant (**Fig. 5B**). **C**: The distribution of $\beta$ coefficients for logistic regression models generated for growth rate and BGC class presence following two resampling approaches. Points represent the mean estimate across all 10,000 bootstrapping replicates and line segments indicate the standard deviation across replicates. Only the 29 BGC classes that were initially significantly correlated with growth rate (**Fig. 5B**) are shown.

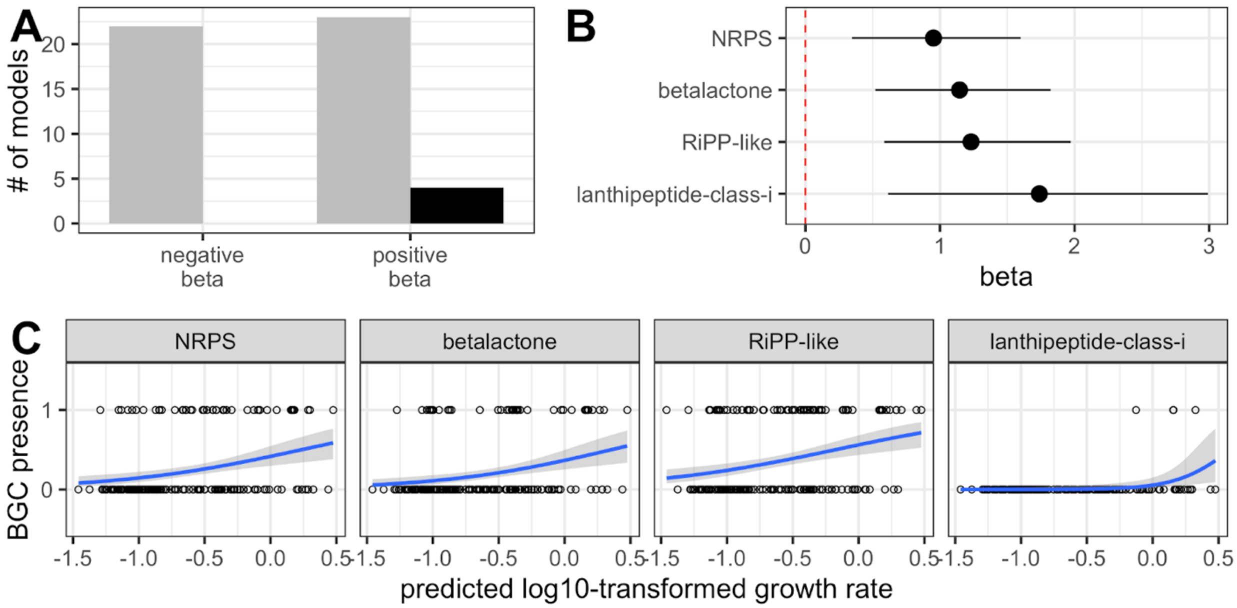


**Supplemental Figure 3. In the Vieira-Silva and Rocha dataset, three of five biosynthetic gene clusters are also predicted to be correlated with growth rate when gRodon-predicted growth rates are used for model generation. A:** Number of logistic regression models with negative and positive $\beta$ values when gRodon-predicted maximum growth rate values are used in place of observed growth rates. The number of models with significant correlations (following Benjamini-Hochberg correction) are colored in black. **B:** $\beta$ coefficients for logistic regression models generated for growth rate and BGC class presence. Line segments indicate 95%

confidence intervals for model estimates. All BGC classes can potentially encode toxins. Three of them were also significantly correlated wth growth rate when observed maximum growth rate was used - betalactones, NRPS, and lanthipeptide class I BGCs – while RiPP-like clusters were not (**Fig. 4B**). Ectoines and siderophores were significantly correlated using observed growth rate but not predicted growth rate. The use of predicted growth rates also increased the value of $\beta$ coefficients for the BGC classes that were significant in both cases, underscoring the differences in both the underlying distribution of the data and the resulting probability distribution. **C:** Regression plots of the four BGC classes with a significant correlation, in order of increasing $\beta$ coefficient. Growth rates are log10-transformed to better observe the range of small values because growth rates across the dataset are left-skewed.

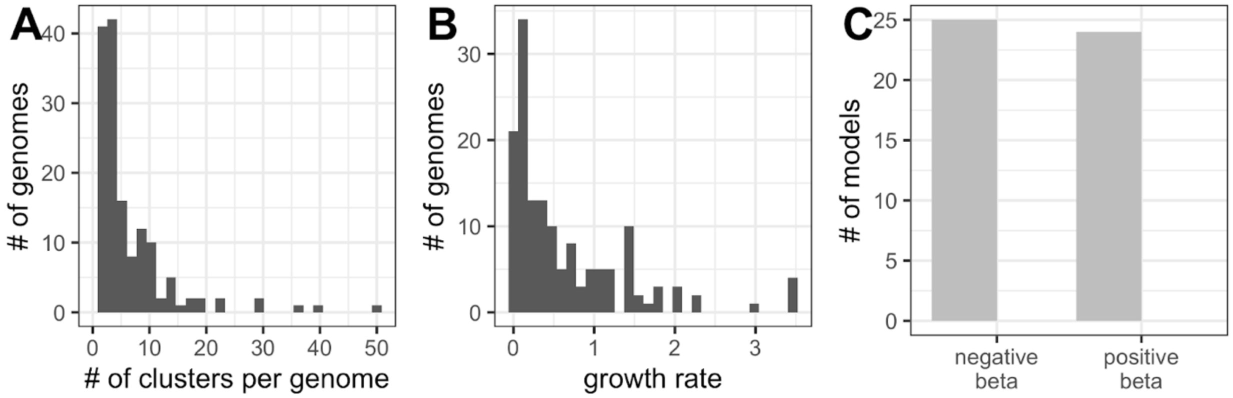


**Supplemental Figure 4. In the Vieira-Silva and Rocha dataset, removing genomes with no detected biosynthetic gene clusters reduces the power of the analysis and eliminates significant correlations between growth rate and presence/absence of certain BGC classes.** **A:** Histogram of total number of detected BGC classes per genome, excluding genomes with no detected BGCs. When genomes without predicted BGCs are excluded, only 148 unique genomes remain in the dataset. The minimum number of BGCs detected in a genome was 1, with a maximum of 50, a mean of 6.5, and a median of 4. **B:** Histogram of growth rates for each dataset, excluding genomes with no detected BGCs. In this reduced dataset, observed growth rates ranged from 0.003 to 3.47 $hr^{-1}$, with a median of 0.35 and a mean of 0.67. **C:** Number of logistic regression models with negative and positive $\beta$ values. The number of models with significant correlations (following Benjamini-Hochberg correction) are colored in black, though there are none due to the reduced power of the analysis.
