## Supplemental File 2 for "The strength of selection for costly toxin production increases with growth rate"

To increase the breadth of our comparative genomics analysis, we used the ncbi_datasets command line tool (ncbi_datasets v. 16.24.1) in August 2024 to download reference genomes for all species represented in the 2023 Danov et al. paper "Toxinome–the bacterial protein toxin database" [1]. The Toxinome dataset is composed of toxin sequences identified in 59,475 genomes across an estimated 8,000 species. We downloaded reference genomes (fna files) for all species for which an annotated reference genome was available on GenBank, leaving us with a dataset of 5,741 species represented by 37,028 unique genomes. GenBank fna files were used as input into gRodon v. 2.0.0 [2]. We aggregated a final dataset by associating individual bacterial strains with the predicted growth rate of its reference genome. For example, 5,014 *Staphylococcus aureus* genomes are represented in the Toxinome dataset, each with a different suite of putative toxin genes. For each of those 5,014 genomes, we assumed that the gRodon-predicted doubling time of the NCBI *S. aureus* reference genome provided a reasonable estimation of growth rate for all individual strains. Lastly, we cleaned the Toxinome dataset to limit it to toxin genes that are most likely to be used in interference competition by removing products corresponding to plasmid maintenance or toxin-antitoxin systems. Our final dataset consisted of IMG genome IDs, species names, reference-genome gRodon-predicted growth rates, and toxin gene product names.

To investigate whether there was a correlation between growth rate and toxin maintenance, we calculated the number of toxin genes identified per genome, as well as the total number of unique types of toxin genes per genome (using product name as type). Generating linear regression models using these data, we found that there was no association between the number of toxin genes in a genome and growth rate (estimate = 0.049), but there was a weak positive correlation between the number of unique types of toxin genes in a genome and growth rate (estimate = 0.8279). We calculated 95% confidence intervals for these models using 10,000 bootstrapping replicates and found that the correlation between unique types and growth rate was always weakly positive, while the correlation between total genes and growth rate was not. We expected that the number of unique types of toxin genes may be a better overall representation of selection for toxin maintenance, indicating the diversity of interference competition systems necessary for a genome's success, while the total number of individual toxin genes should be more likely to be confounded by species-specific rates of genomic evolution. As a result, we posited that this dataset was weakly supportive of our hypothesis, especially given the enormous size and taxonomic diversity of the Toxinome database.

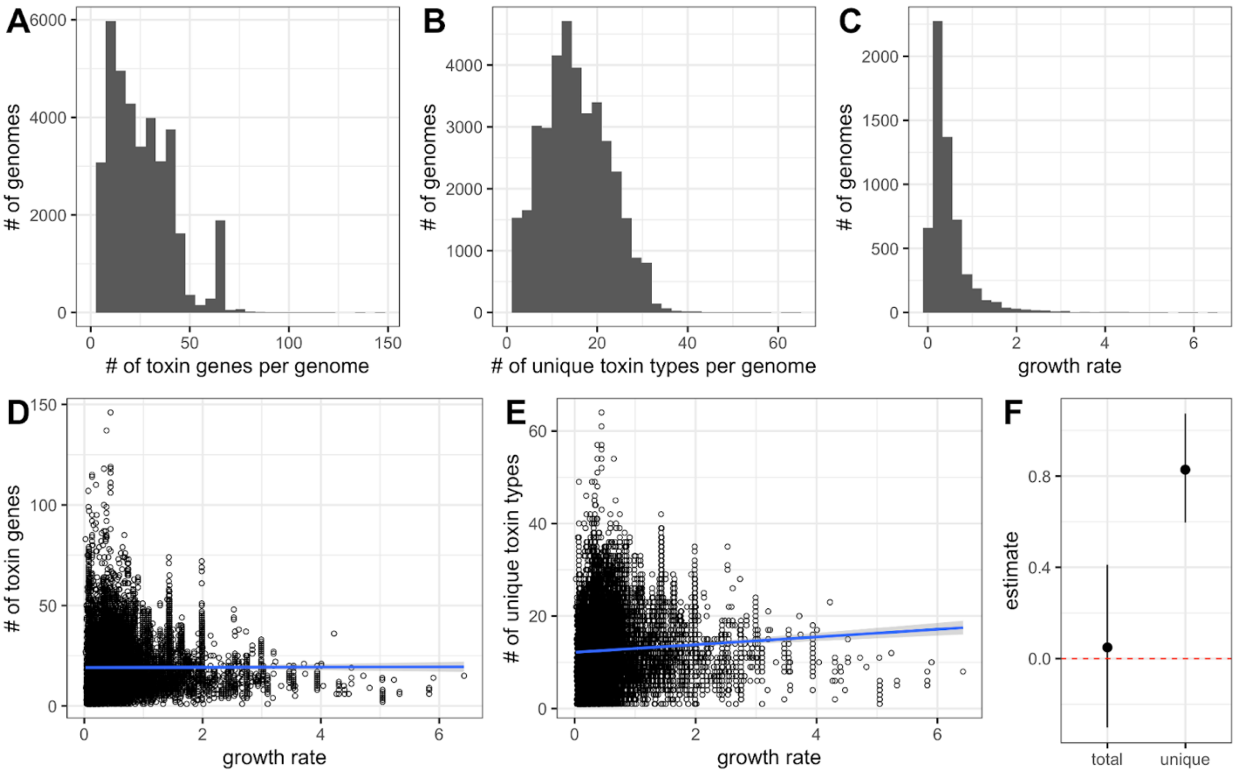


**Supplemental Figure 5. In a large dataset of predicted toxin genes, the number of unique types of toxin genes present in a genome is weakly correlated with growth rate. A:**

Histogram of total number of predicted toxin genes per genome (min: 1, max: 146, mean: 25.14, median: 22). **B:** Histogram of number of unique types of predicted toxin genes per genome (min: 1, max: 64, mean: 15, median: 15). **C:** Histogram of predicted growth rates across all species represented (min: 0.003, max: 6.42, mean: 0.47, median: 0.33). **D:** Linear regression model between growth rate and number of toxin genes present in a genome, with an estimate of 0.049 and an R-squared of 4.4e-6. **E:** Linear regression model between growth rate and number of unique types of toxin genes present in a genome, with an estimate of 0.8279 and an R-squared of 0.004. **F:** 95% confidence interval of linear regression estimates based on 10,000 bootstrapping replicates. The dotted red line represents the switch from a positive to a negative correlation.
